# *Birnaviridae* virus factories show features of liquid-liquid phase separation, and are distinct from paracrystalline arrays of virions observed by electron microscopy

**DOI:** 10.1101/2021.11.16.468875

**Authors:** Vishwanatha R. A. P. Reddy, Elle A. Campbell, Joanna Wells, Jennifer Simpson, Salik Nazki, Philippa C. Hawes, Andrew J. Broadbent

## Abstract

To gain more information about the nature of *Birnaviridae* virus factories (VFs), we used a recombinant infectious bursal disease virus (IBDV) expressing split-GFP11 tagged to the polymerase (VP1) that we have previously shown is a marker for VFs in infected cells expressing GFP1-10. We found that VFs co-localized with 5-ethynyl uridine in the presence of actinomycin D, confirming they were the site of *de novo* RNA synthesis, and VFs were visible in infected cells that were fixed and permeabilized with digitonin, demonstrating that they were not membrane bound. Fluorescence recovery after photobleaching (FRAP) a region of interest within the VFs occurred rapidly, recovering from approximately 25% to 87% the original intensity over 146 seconds, and VFs were dissolved by 1,6-hexanediol treatment, demonstrating they showed properties consistent with liquid-liquid phase separation. There was a lower co-localization of the VF GFP signal with the capsid protein VP2 (Manders’ coefficient (MC) 0.6), compared to VP3 (MC, 0.9), which prompted us to investigate the VF ultrastructure by transmission electron microscopy (TEM). In infected cells, paracrystalline arrays (PAs) of virions were observed in the cytoplasm, as well as discrete electron dense regions. Using correlative light and electron microscopy (CLEM), we observed that the electron dense regions correlated with the GFP signal of the VFs, which were distinct from the PAs. In summary, *Birnaviridae* VFs are sites of *de novo* RNA synthesis, are not bound by a membrane, show properties consistent with liquid-liquid phase separation, and are distinct from the PAs observed by TEM.

**Importance:** Members of the *Birnaviridae* infect birds, fish and insects, and are responsible for diseases of significant economic importance to the poultry industry and aquaculture. Despite their importance, how they replicate in cells remains poorly understood. Here, we show that the *Birnaviridae* virus factories are not membrane bound, demonstrate properties consistent with liquid-liquid phase separation, and are distinct from the paracrystalline arrays of virions observed by transmission electron microscopy, enhancing our fundamental knowledge of virus replication that could be used to develop strategies to control disease, or optimize their therapeutic application.

## Introduction

The *Birnaviridae* family is divided into 4 genera that infect birds (*Avibirnavirus*), fish (*Aquabirnavirus* and *Blosnavirus*), and insects (*Entomobirnavirus*) (1). Within the genus *Avibirnavirus*, infectious bursal disease virus (IBDV) is of major concern to the global poultry industry due to morbidity, mortality, and immunosuppression in chickens that lead to an increase in secondary infections (2). IBDV is controlled through vaccination (3), and recombinant vaccine strains also show promise as vectors for the expression of antigens to control other infections (4, 5). The *Avibirnavirus,* chicken proventricular necrosis virus (CPNV), has also recently been proposed to be the cause of transmissible viral proventriculitis (TVP)(6). Within the genus *Aquabirnavirus*, infectious pancreatic necrosis virus (IPNV) infects salmonid fish, for example salmon and trout, causing high mortality in fry and fingerlings, and a number of related viruses are associated with disease in a wide variety of fish, mollusks, and crustaceans, resulting in significant economic losses to aquaculture (1, 7). In addition, Blotched Snakehead virus, another birnavirus infecting fish, is the sole member of the genus *Blosnavirus*, and is more closely related to the *avibirnaviruses* than the *aquabirnaviruses* (8). Members of the genus *Entomobirnavirus* infect insects, and include Drosophila X virus (DXV) which has played an important role in the study of innate immunity and siRNA pathways (9, 10), Culex Y virus (CYV), and Port Bolivar virus that have been identified in mosquito cells (11, 12).

Despite their abundance and importance, many aspects of the replication of the *Birnaviridae* remain poorly understood. As the viruses are non-enveloped, and contain a segmented double stranded (ds)RNA genome, assumptions of their replication strategies have been extrapolated from knowledge of more well-studied dsRNA viruses, for example the *Reoviridae* family. However, significant differences between the strategies used by the *Birnaviridae* and the *Reoviridae* are beginning to be uncovered. For example, birnaviruses do not make use of a core particle during the replication cycle (13). Instead, they possess a single capsid layer and uncoat in the endosome, whereupon the genome, combined in a ribonucleoprotein (RNP) complex with the viral polymerase (VP1) and the nonstructural protein VP3, is extruded into the cytoplasm (14). In the cytoplasm, the VP3 tethers the RNP complex to the cytoplasmic leaflet of endosomal membranes (15) via interacting with phosphatidylinositol-3-phosphate (16), and it is thought that the virus co-opts these cellular membranes to seed the formation of a replication complex, or virus factory (VF), to provide a microenvironment more conducive to replication.

In order to provide more information on the nature of the *Birnaviridae* VFs, we previously rescued a replication-competent IBDV, strain PBG98, containing a split green fluorescent protein (GFP)11 tag fused to the C-terminus of the VP1 polymerase (PBG98-VP1-GFP11) (17). In infected cells expressing GFP1-10, the GFP11 tag complemented the GFP1-10 molecule and the resultant complete GFP molecule fluoresced green. The GFP signal in PBG-VP1-GFP11-infected cells co-localized with other components of the RNP complex, VP3, and dsRNA, visualized by immunofluorescence microscopy, and with genome segments A and B, visualized by fluorescence in situ hybridization (FISH). We therefore concluded it was a marker for VFs. Using live cell imaging, we found that the VFs moved in the cytoplasm and coalesced together over time (17). Here, we build on this work to further understand the nature of the *Birnaviridae* VFs, investigating the possibility they are formed from liquid-liquid phase separation using live cell imaging approaches, and defining their ultrastructure by transmission electron microcopy (TEM), and correlative light and electron microscopy (CLEM).

## Materials and Methods

### Cell-lines and antibodies

DF-1 cells (chicken embryonic fibroblast cells, ATCC number CRL-12203) were maintained in Dulbecco’s modified Eagle’s medium (DMEM) (Sigma-Aldrich, Merck, UK), supplemented with 10% heat inactivated fetal bovine serum (hiFBS) (Gibco, Thermo Fisher Scientific, UK). The primary antibodies used in this study were raised against tubulin (SantaCruz Biotechnology, UK), PDI (Enzo Life Science, Farmingdale, NY), VP2, and VP3 (18). Primary antibodies were diluted 1:100 and secondary antibodies conjugated to Alexa flour 488-or 568 (Invitrogen, Thermo Fisher Scientific, UK) were diluted 1:500 in 4% bovine serum albumin (BSA) (Sigma-Aldrich, Merck, UK).

### Viruses

The cell-culture adapted Avian Reovirus (ARV) strain, S1133, was obtained from Charles River (Wilmington, MA, USA), and a molecular clone of the cell-culture adapted IBDV strain, PBG98, was rescued as previously described (17). Three alanine residues were added to the N terminus of the GFP11 tag as a linker and a stop codon was added to the C terminus to make the amino acid sequence: AAARDHMVLHEYVNAAGIT-Stop. The nucleotide sequence encoding this was added to IBDV segment B, which encodes VP1, prior to the 3’ non-coding region, and the recombinant virus, PBG98-VP1-GFP11, was rescued as previously described (17). Stocks of the viruses were grown in DF-1 cells and titrated in 96-well plates (Falcon, Corning, UK) seeded with DF-1 cells at a density of 4 × 10^4^ cells per well. Wells were infected in quadruplicate with a 10-fold dilution series of supernatant. After 5 days, wells were inspected for signs of cytopathic effect (cpe) and the titer of virus determined as per the method by Reed and Muench (19), and expressed as Tissue Culture Infective Dose-50 (TCID50).

### Immunofluorescence Microscopy

DF-1 cells were seeded onto coverslips (TAAB, UK) in 24-well plates (Falcon, Corning, UK) at a density of 1.6 ×10^5^ per well and transfected with GFP1-10 using lipofectamine 2000, 24 hours prior to infection with the PBG98-VP1-GFP11 virus. At the relevant time-points, cells were fixed with a 4% paraformaldehyde solution (Sigma-Aldrich, Merck, UK) for 20 minutes, permeabilized, and blocked with a 4% BSA solution for 30 minutes at room temperature. Cells were permeabilized with a solution of 0.1% Triton X-100 (Sigma-Aldrich, Merck, UK) for 15 minutes, unless otherwise stated. In order to evaluate whether VFs were membrane-bound, cells were permeabilized with 1:2,000 digitonin (Caltag-Medsystems, Buckingham, UK) for 30 minutes. Cells were then incubated with the appropriate primary antibody for 1 hour, washed with PBS, and incubated with the secondary antibody for 1 hour in the dark at room temperature. Cells were again washed, incubated in a solution of 4,6-diamidino-2-phenylindole dihydrochloride (DAPI, Invitrogen Thermo Fisher Scientific, UK), washed in water, mounted with Vectashield (Vector Laboratories Inc, CA), and imaged with a Leica TCS SP5 confocal microscope.

### Visualizing 5-Ethynyl Uridine (EU) signal

DF-1 cells were transfected with GF1-10 and infected with the PBG98-VP1-GFP11 virus 24 hours post-transfection (hpt) at an MOI of 1. Thirteen hours post-infection (hpi), cells were treated with 10μg/mL of actinomycin D to prevent cellular transcription for 4 hours, after which 1mM of EU was added to the cells that became incorporated into newly synthesized RNA molecules. 1 hour after EU treatment, cells were fixed, permeabilized with 0.1% triton-X, and stained with a Click-iT RNA imaging kit (Thermo Fisher Scientific) to visualize the EU, according to the manufacturer’s instructions. Briefly, permeabilized cells were incubated in Click-iT reaction cocktail containing Alexa-Fluor 594 azide for 30 minutes at room temperature, washed with Click-iT reaction rinse buffer, and stained with DAPI. Cells were washed in water, mounted in Vectashield and imaged with a Leica TCS SP5 confocal microscope.

### Live cell imaging

DF-1 cells were seeded into a Chambered #1.0 Borosilicate Coverglass slide (Nunc, Lab-Tek, Sigma-Aldrich, Merck, UK) at a density of 8 × 10^4^ per well and cultured in 1ml of DMEM supplemented with 10% hiFBS. Twenty four hours after seeding, cells were transfected with GFP1-10 and then infected with the PBG98-VP1-GFP11 virus 24 hpt. Sixteen hpi, cells were incubated in Leibovitz’s L-15 media without phenol red (Gibco, Thermo Fisher Scientific, UK), and imaged live with a Leica TCS SP8 confocal microscope in a climate controlled chamber. Cells were either treated with a solution of 4% 1,6-hexanediol (Sigma-Aldrich) diluted in L-15 media, or were mock-treated with L-15 media alone. For each time course, 50 z-stacks were imaged in total, with each stack being acquired at an interval of 20 seconds.

For fluorescence recovery after photobleaching (FRAP), one region of interest (ROI) with diameter 0.6μm was photobleached within infected cells, inside a VF. A region adjacent to the factory (with no detectable fluorescence) was also imaged as a control. For each dataset, 5 images were taken pre-bleach, followed by 1 bleach scan (488nm laser at 70%), then 170 post-bleach images were acquired over the course of the experiment (146 seconds) to monitor fluorescence recovery. Scans with high levels of drift were excluded from further analysis. Recovery datasets were also collected from non-bleached ROIs in the same cells as controls.

### Confocal Imaging Analysis

An overlap coefficient (an alternative to the Pearson’s correlation coefficient created by Manders *et al*) was used as the main statistical test to describe co-localization using Image J software (National Institutes of Health, NIH). Images were only considered in the analysis if they had a Coste’s significant level of 0.95 or above (20). For FRAP, Fluorescence intensity datasets were exported from the LAS X software, normalized for background and acquisition photobleaching using the easyFRAP web tool (21) and fit to a recovery curve using ImageJ.

### Transmission Electron Microscopy (TEM)

DF-1 cells were seeded onto 13mm Thermanox coverslips (Thermo Fisher Scientific, UK) and infected with either IBDV strain PBG98, PBG98-VP1-GFP11, or ARV strain S1133 at an MOI of 1. At 10, 18 and 20 hpi, cells were fixed in phosphate buffered 2% glutaraldehyde (Agar Scientific) for 1 hour before being fixed for a further hour in aqueous 1% osmium tetroxide (Agar Scientific). The samples were dehydrated in an ethanol series; 70% for 30 min, 90% for 15 min and 100% three times for 10 min each. A transitional step of 10 min in propylene oxide (Agar Scientific) was undertaken before the samples were infiltrated with 50:50 mix of propylene oxide and epoxy resin (Agar Scientific) for 1 hour. After a final infiltration of 100% epoxy resin for 1 hour, the samples were embedded and polymerized overnight at 60°C. 80nm thin sections were cut, collected onto copper grids (Agar Scientific) and grid stained using Leica EM AC20 before being imaged at 100kV in a FEI Tecnai 12 TEM with a TVIPS F214 digital camera.

### Electron Tomography (ET)

DF-1 cells were seeded onto 13mm Thermanox coverslips (Thermo Fisher Scientific, UK) and infected with IBDV strain PBG98 at an MOI of 1. At 18 hpi, cells were prepared for TEM. 250nm sections were cut and collected onto Formvar coated copper hex50 grids (Agar Scientific) and 10μl of 10nm gold antibody (BBI Solutions) was placed on the grids to act as fiducial markers. Grids were imaged at 200kV in a JEOL 2100F TEM with a TVIPS F416 digital camera. Using Serial EM, data sets were collected every 1^⍛^ over a 120^⍛^ tilt series. Reconstruction and modelling of the data sets was carried out using IMOD.

### Correlative Light and Electron Microscopy (CLEM)

DF-1 cells were seeded onto gridded glass bottom dishes (MatTek) at a density of 8 × 10^4^ per well and cultured in 1mL of DMEM supplemented with 10% hiFBS. Twenty four hours after seeding, cells were transfected with GFP1-10 and then infected with the PBG98-VP1-GFP11 virus 24 hpt. Eighteen hpi, cells were stained with Hoechst 33342 (Thermo Fisher Scientific) and selected grid squares were imaged live in L-15 media with a Leica TCS SP8 confocal microscope in a climate controlled chamber. The samples were fixed in phosphate buffered 2% glutaraldehyde (Agar Scientific) for 1 hour before being fixed for a further hour in aqueous 1% osmium tetroxide (Agar Scientific). The samples were dehydrated in an ethanol series; 70% for 30 min, 90% for 15 min and 100% three times for 10 min each. After infiltration of 100% epoxy resin (Agar Scientific) for 2 hours, the samples were embedded and polymerized overnight at 60°C. The glass coverslips were removed with liquid nitrogen and the appropriate grid squares located. 80nm thin sections were cut, collected onto copper grids (Agar Scientific) and grid stained using Leica EM AC20. The specific cells imaged in the confocal were located and imaged at 100kV in a FEI Tecnai 12 TEM with a TVIPS F214 digital camera.

## Results

### IBDV VFs were the sites of *de novo* RNA synthesis

DF-1 cells were transfected with GFP1-10 and infected with the PBG98-VP1-GFP11 virus. Infected cells were treated with actinomycin D to prevent cellular transcription, after which EU was added to the cells that became incorporated into newly synthesized mRNA molecules. After EU treatment, cells were stained with a Click-iT RNA imaging kit to visualize EU, and stained with DAPI (Figure 1). In uninfected control cells lacking actinomycin D treatment, the EU signal, representing cellular mRNAs, was visible in the nucleus, as expected given that transcription is intranuclear. This intranuclear signal was lost in cells treated with actinomycin D, which blocked cellular transcription. VFs were visible in infected cells that were not treated with actinomycin D or EU (Figure 1A), and in cells that were treated with actinomycin D, but not EU (Figure 1B), indicating that blocking cellular transcription at this time point did not prevent VF formation. Infected cells that were not treated with actinomycin D, and were stained with EU had an EU signal in the nucleus, likely representative of cellular mRNA synthesis, and a signal that co-localized with the VFs, indicating that the PBG98-VP1-GFP11 VFs were also sites of *de novo* RNA synthesis (Figure 1C). In infected cells treated with actinomycin D, the only EU signal detected co-localized with the VFs in the cytoplasm (Figure 1D), indicating that the RNA synthesis that took place in the VFs was independent of the cellular machinery, consistent with the action of the viral polymerase.

**Figure 1.**
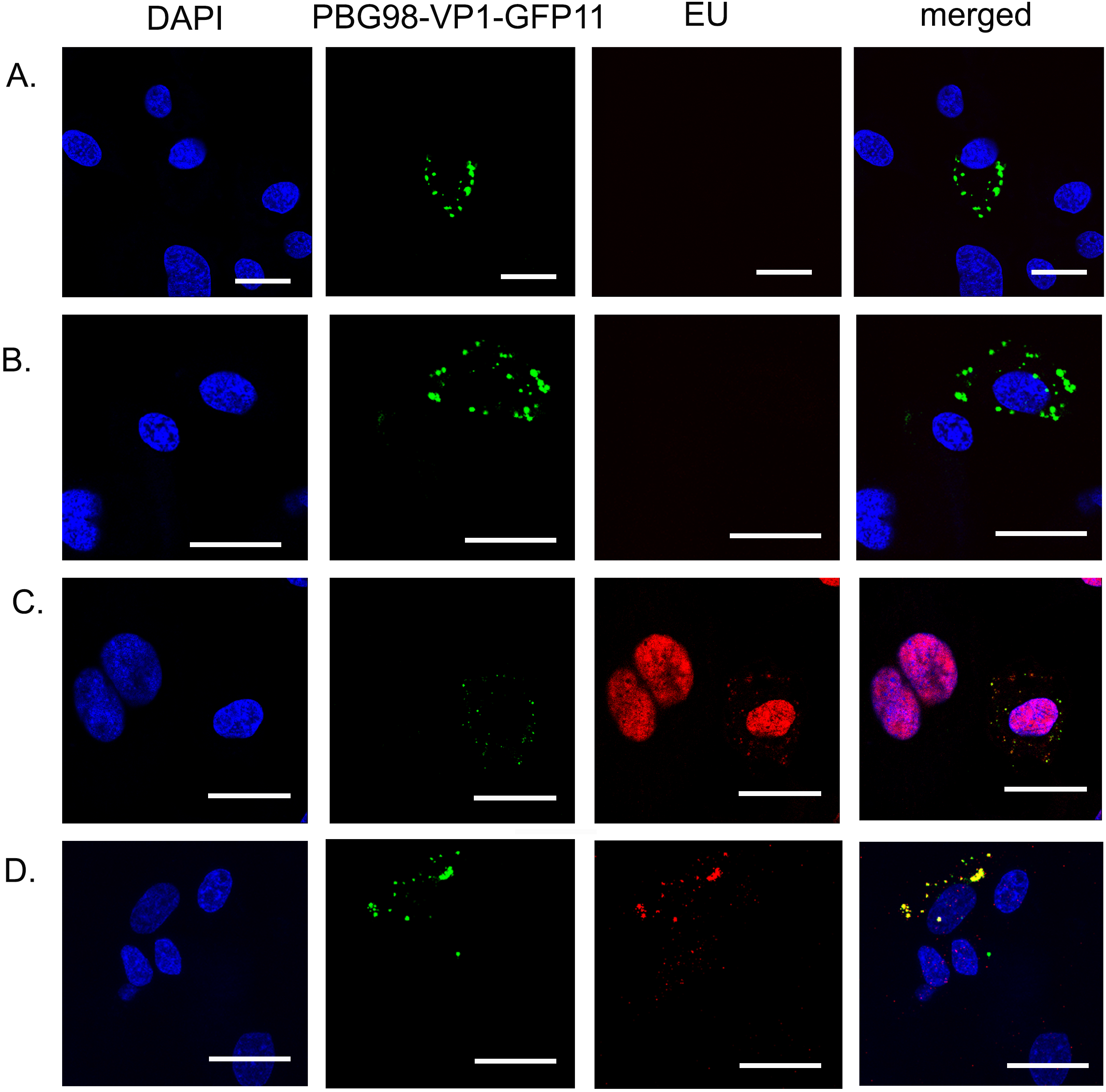
IBDV VFs were the sites of *de novo* RNA synthesis. DF-1 cells were transfected with GFP1-10 and infected with the PBG98-VP1-GFP11 virus 24hpt at an MOI of 1 (A-D). Thirteen hpi, cells were either treated with actinomycin D for 4 hours to prevent cellular transcription (B and D), or mock treated with media (A and C), after which cell cultures were either treated with 1mM EU that became incorporated into newly synthesized RNA molecules (C and D), or mock treated with media alone (A and B). One hour after EU treatment, all cells were stained with a Click-iT RNA imaging kit to visualize EU, and stained with DAPI. Scale bars, 20μm.

### IBDV VFs were not bound by a membrane

DF-1 cells were transfected with GFP1-10 and infected with the PBG98-VP1-GFP11 virus. Infected cells were fixed and permeabilized with either Triton X-100, which permeabilized the plasma membrane and internal membranes, or digitonin, which permeabilized the plasma membrane but not intracellular membranes. Cells were stained with primary antibodies raised against tubulin (a cytoplasmic marker), PDI (present within the endoplasmic reticulum and therefore a marker of intracellular membrane permeabilization), or VP3, followed by secondary antibodies conjugated to Alexafluor 568 (Figure 2). As expected, the cytoplasmic tubulin was visible following either permeabilization method, but PDI was not visible when cells were permeabilized with digitonin as it was contained within a membraned organelle (Figure 2C and D). The PBG98-VP1-GFP11 signal was present in cells permeabilized with either triton X or digitonin, which co-localized with VP3, demonstrating that the VFs were not bound by a membrane.

**Figure 2.**
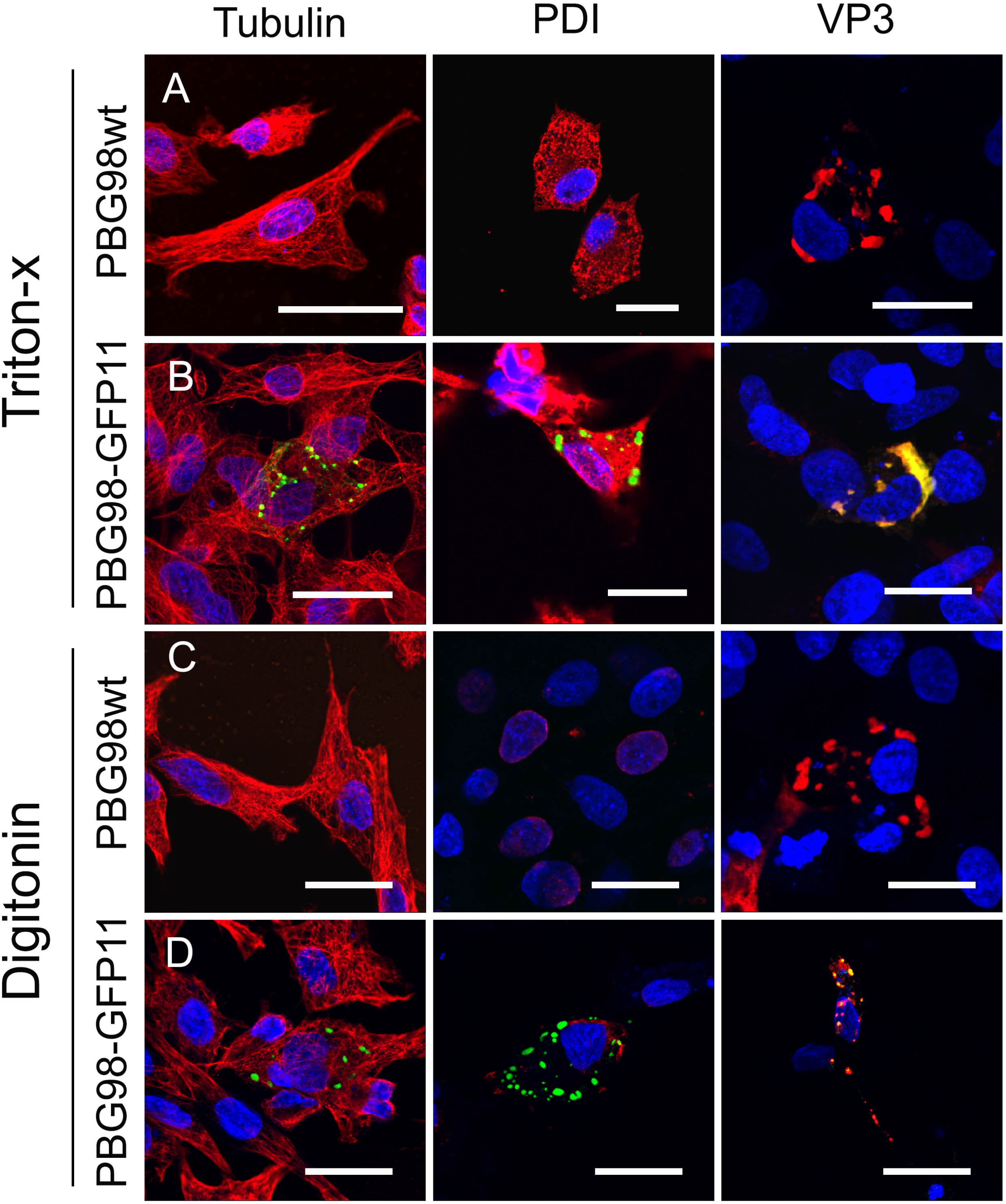
IBDV VFs were not bound by a membrane. DF-1 cells were either infected with recombinant PBG98 wt (A and C), or transfected with GFP1-10 and infected with the PBG98-VP1-GFP11 virus (B and D). Infected cells were fixed and permeabilized with either Triton X-100 (A and B), or digitonin (C and D). Cells were then stained with primary antibodies raised against tubulin, PDI, or IBDV VP3, followed by secondary antibodies conjugated to Alexafluor 568, and DAPI. Scale bars, 20 μm.

### IBDV VFs had properties consistent with liquid-liquid phase separation

DF-1 cells were transfected with GFP1-10 and infected with the PBG98-VP1-GFP11 virus. ROIs within the VFs were photobleached to approximately 25% of their original intensity, and fluorescence recovery was determined over time. The VFs showed a rapid recovery in fluorescence, to approximately 87% (standard deviation (SD) 3.5) of the original intensity during the course of the experiment (146 seconds) (Figure 3A). In a separate experiment, DF-1 cells were transfected with GFP1-10 and infected with the PBG98-VP1-GFP11 virus. Infected cells were treated with 4% 1,6-hexanediol, which has been shown to dissolve the VFs of rotavirus (22). In the presence of 1,6-hexanediol, the VFs of PBG98-VP1-GFP11 dissolved (Figure 3B and Movie S2), whereas the VFs of mock-treated cells were visible throughout the experiment (Figure 3B and Movie S1). These data, taken together with our previous study showing VFs rapidly moved in the cytoplasm and coalesced together (17), demonstrated that IBDV VFs had properties consistent with liquid-liquid phase separation.

**Figure 3.**
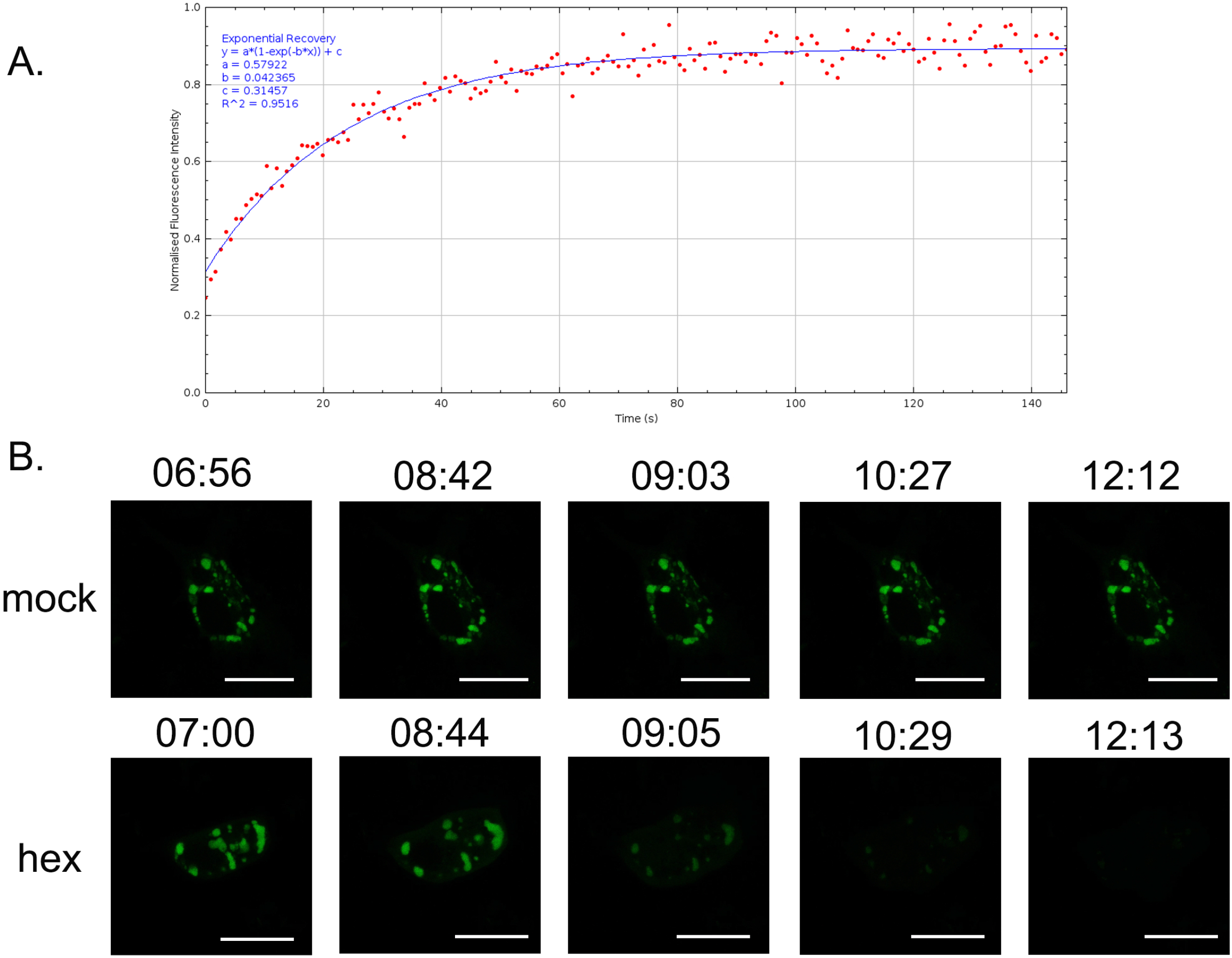
IBDV VFs had properties consistent with liquid-liquid phase separation. DF-1 cells were transfected with GFP1-10 and then infected with the PBG98-VP1-GFP11 virus 24 hpt. At 16 hpi, one ROI with a 0.6μm diameter was photobleached inside a VF to approximately 25% the original intensity with the 488nm Argon laser. Then, 170 post-bleach images were acquired during the experiment (146 seconds). The fluorescence intensity was normalized to the background, and plotted against time (A). Infected cells were either mock-treated with media, or were treated with a solution of 4% 1,6-hexanediol (hex) 16hpi and imaged live over a time course of 17 minutes. For each time course, 50 z-stacks were imaged in total, with each stack being acquired at an interval of 20 seconds. Images of different time points within the time course (indicated by the time stamp (minutes: seconds) are shown (B). Scale bars, 20 μm.

### Virus Factory like structures (VFLs) formed in the cytoplasm of cells transfected with VP3-GFP, but not other viral proteins

DF-1 cells were transfected individually with plasmids encoding each of the viral proteins VP1, 2, 3, 4 and 5, tagged to GFP, and imaged by confocal microscopy. Cells transfected with either GFP-VP1 or VP2-GFP had a fluorescence signature that was present throughout the cytoplasm, whereas cells transfected with GFP-VP3 showed puncta in the cytoplasm reminiscent of the virus factories. We termed these virus factory-like (VFL) structures. Cells transfected with GFP-VP4 contained wave-shaped structures in the cytoplasm, and cells transfected with GFP-VP5 had a signal that appeared to be near or at the plasma membrane (Figure 4 A). When a small ROI of a VP3-GFP VFL was photobleached to approximately 20% the original intensity, the fluorescence recovered to approximately 46% the original intensity (SD 1.24, n = 11) during the course of the experiment. In contrast, when a small ROI of a VP4-GFP structure was photobleached, there was no increase in intensity, which remained at approximately 22% the original value (SD 0.73, n = 9) throughout the experiment (Figure 4B). Taken together, these data demonstrate that VFLs were observed in cells transfected with VP3-GFP alone, but not the other proteins, and that VP3 VFLs showed some liquid-like properties.

**Figure 4.**
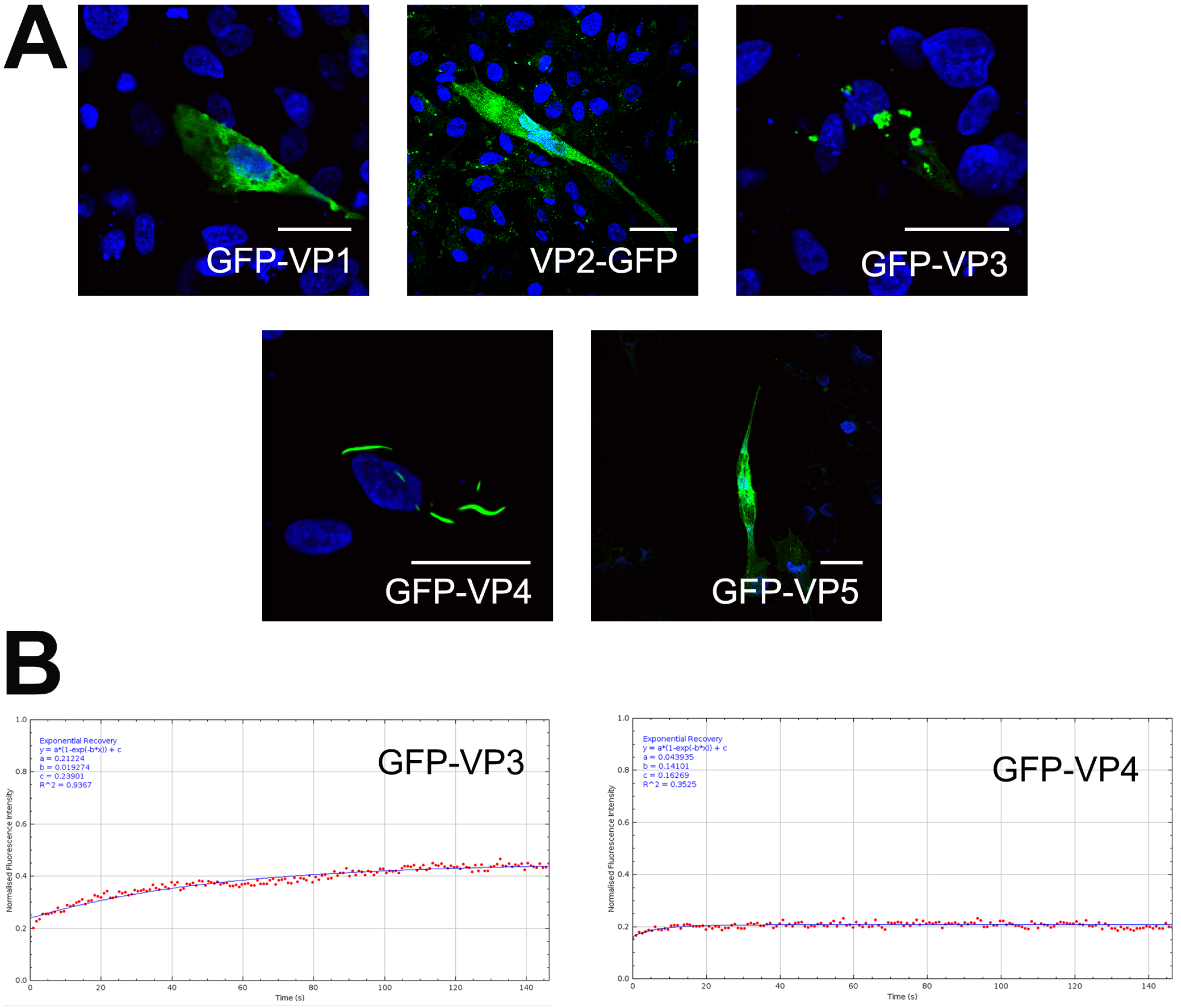
Virus Factory like structures (VFLs) formed in the cytoplasm of cells transfected with VP3-GFP, but not other viral proteins. DF-1 cells were transfected individually with plasmids encoding each of the viral proteins VP1-5, tagged to GFP. Transfected cells were fixed 24hpt, stained with DAPI, and imaged. Scale bars, 20 μm (A). DF-1 cells transfected with GFP-VP3 and GFP-VP4 were imaged live and, at 16 hpt, one ROI within one recorded cell was photobleached to approximately 20% the original intensity with the 488nm Argon laser. Then, 170 post-bleach images were acquired during the experiment (146 seconds). The fluorescence intensity was normalized to the background, and plotted against time (B).

### VFs had a lower co-localization with VP2 than VP3 by confocal microscopy

In a previous study, we demonstrated that the VFs had a high co-localization with VP3. In this experiment, we observed the co-localization of the VF GFP signal with VP2, VP3, and VP4 (Figure 5A), and quantified this using a Manders’ coefficient (MC) (Figure 5B). We found the MC to be 0.9 for VP3, and 0.2 for VP4, consistent with strong and weak co-localization, respectively. In addition, we observed a lower co-localization with the capsid protein, VP2 (MC, 0.6) than VP3, that prompted us to investigate the ultrastructure of the VF by TEM.

**Figure 5.**
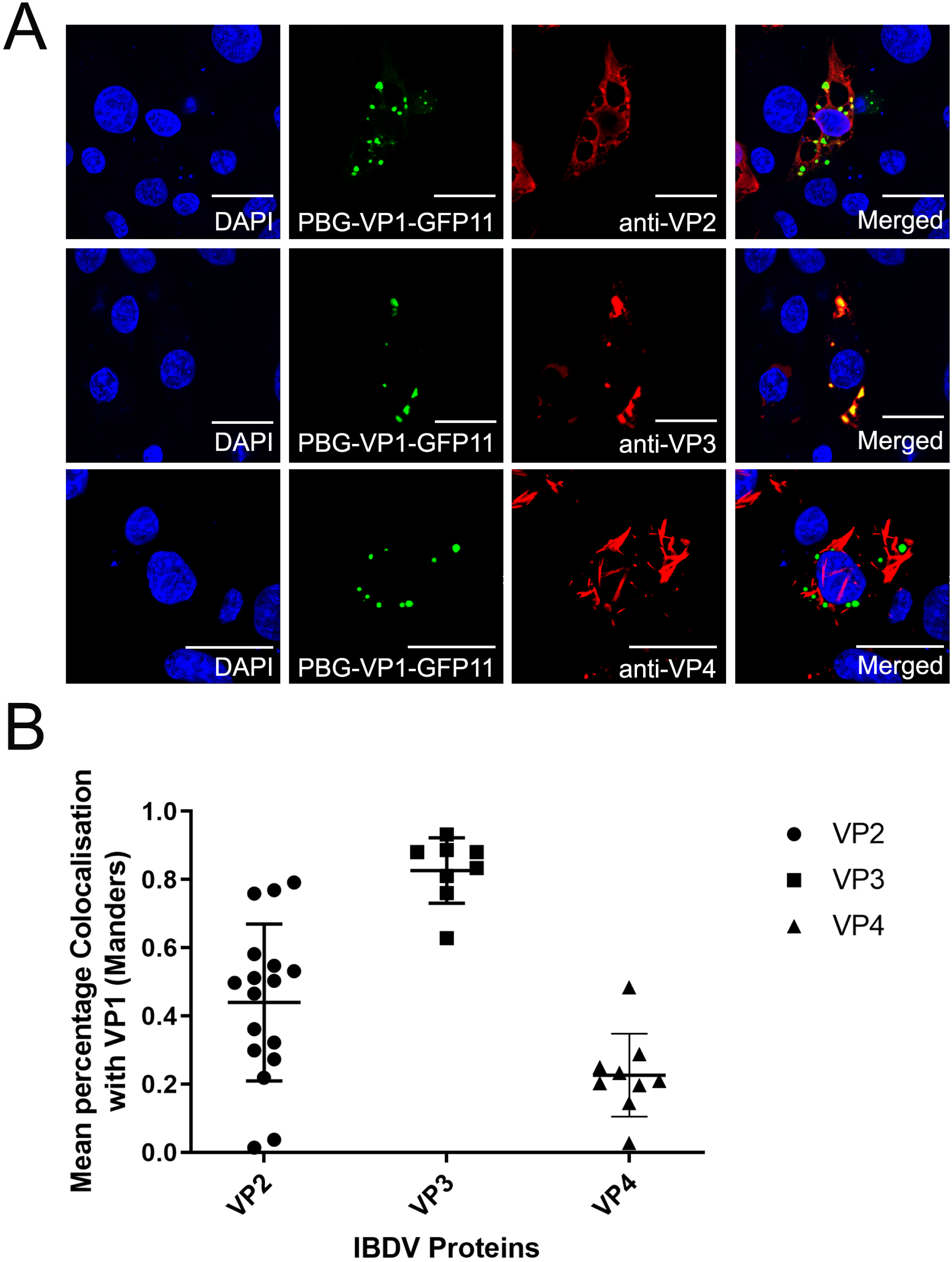
VFs had a lower co-localization with VP2 than VP3 by confocal microscopy. DF-1 cells were transfected with GFP1-10 and infected with the PBG98-VP1-GFP11 virus 24hpt at an MOI of 1. At 24 hpi, infected cells were fixed, permeabilized, and stained with anti-VP2, anti-VP3, or anti-VP4, followed by secondary antibodies conjugated to Alexafluor 568, and DAPI. Scale bars, 20 μm (A). An overlap coefficient, (an alternative to the Pearson’s correlation coefficient, created by Manders et al) was used to quantify the co-localization of the VP1-GFP signal of the VFs with the VP2, VP3, and VP4 signals (B).

### Paracrystalline Arrays (PAs) of virions and electron-dense regions of the cytoplasm were observed in infected cells by TEM

DF-1 cells were either transfected with GFP1-10 and then infected with PBG98-VP1-GFP11 (Figure 6A), or were infected with the avian reovirus (ARV) strain S1133 (Figure 6B), or the recombinant wild-type (wt) PBG98 (Figure 6C), and imaged by TEM. We observed PAs of virions in the cytoplasm of cells infected with IBDV strain PBG98-VP1-GFP11 (Figure 6A). Upon further observation, we were unable to discern any other structures within the PAs, even by electron tomography (ET) (Movie S3). In contrast, DF-1 cells infected with ARV strain S1133 contained VFs that had a more complex ultrastructure, including an electron –dense viroplasm, as well as PAs of virions (Figure 6B). In IBDV-infected cells, we observed discrete electron-dense regions in the cytoplasm that were distinct from the PAs. These were observed in cells transfected with GFP1-10 and infected with PBG98-VP1-GFP11 (data not shown), as well as DF-1 cells that were not expressing GFP1-10 and were infected with wt PBG98 (Figure 6C).

**Figure 6.**
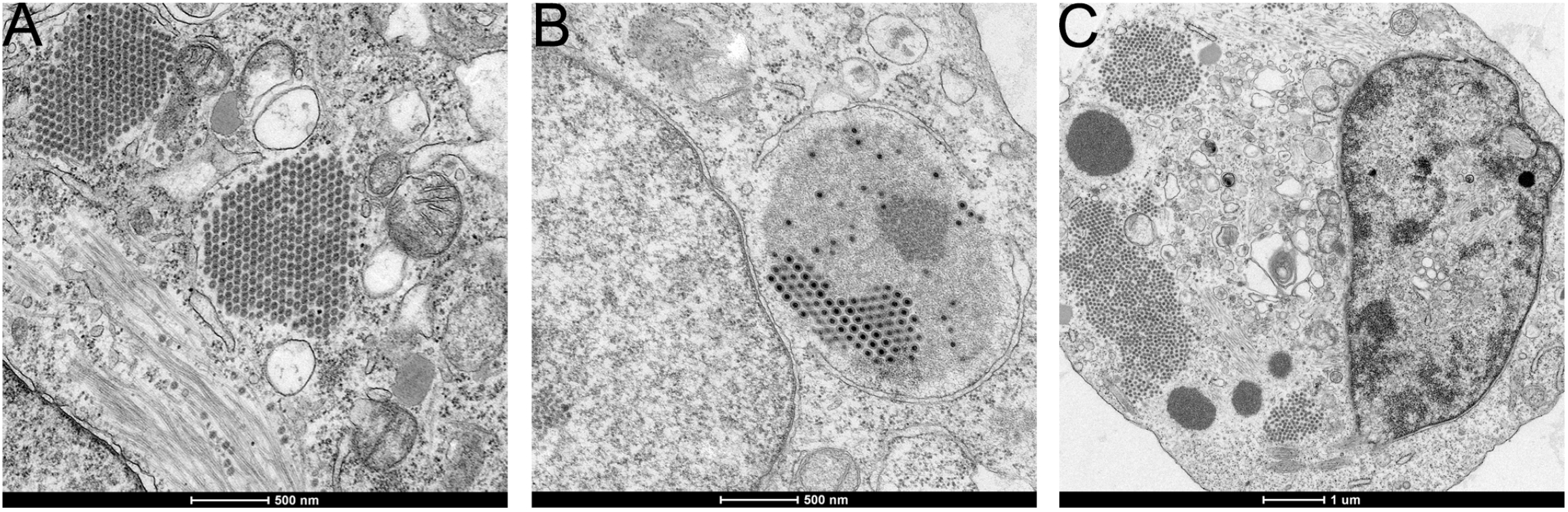
PAs of virions and electron-dense regions of the cytoplasm were observed in infected cells by TEM. DF-1 cells were either transfected with GFP1-10 and infected 24hpt with IBDV strain PBG98-VP1-GFP11 virus (A), or DF-1 cells were infected with the ARV strain S1133 (B), or the recombinant wt IBDV strain PBG98 at an MOI of 1. At 18 (A), 20 (B), or 10 (C) hpi, cells were fixed, prepared, and imaged by TEM.

### VFs correlated with discrete electron-dense regions of the cytoplasm that were distinct from the PAs

In order to determine what structures the VFs we observed by confocal microscopy correlated with by TEM, DF-1 cells were seeded onto gridded glass coverslips, transfected with GFP1-10, and infected with the PBG98-VP1-GFP11 virus. Infected cells were stained with Hoechst 33342 and imaged live by confocal microscopy (Figure 7A and B). After imaging by confocal microscopy, cells were fixed and prepared for TEM. The same cell that was imaged by confocal microscopy was imaged by TEM (Figure 7C), and the two images overlaid (Figure 7 D and E). We observed that the VFs correlated with the electron dense regions of the cytoplasm and were distinct from the PAs of virions (Figure 7).

**Figure 7.**
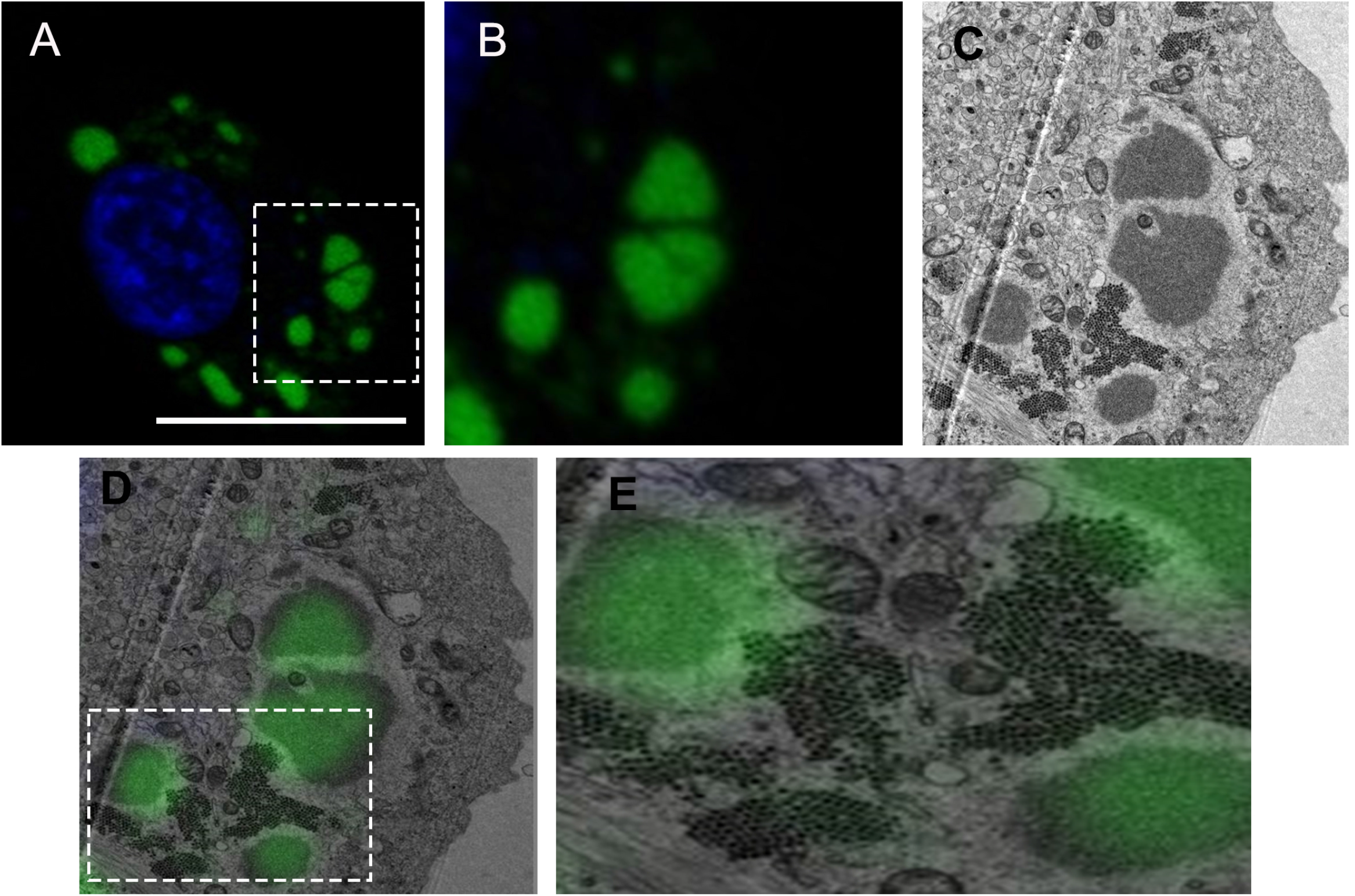
VFs correlated with discrete electron-dense regions of the cytoplasm that were distinct from the PAs. DF-1 cells were seeded onto gridded coverslips, transfected with GFP1-10, and infected with the PBG98-VP1-GFP11 virus 24hpt at an MOI of 1. Eighteen hpi, cells were stained with Hoechst 33342, and cells on selected grid squares were imaged live by confocal microscopy. Scale bar, 20 μm (A). An image of the boxed region is shown enlarged (B). The same cells were then processed and imaged by TEM. The same cell from figure A and B is shown in (C). The confocal image was then overlaid on top of the TEM image and the two correlated (D). An image of the boxed region is shown enlarged (E).

## Discussion

We previously demonstrated that in cells transfected with GFP1-10 and infected with an IBDV with GFP11 fused to the C terminus of VP1, the GFP1-10 and GFP11 molecules came together to form the full-length GFP structure, which fluoresced green, the signal of which co-localized with viral ribonucleoprotein (vRNP) components VP3 and dsRNA (17). In the present study, we extended these observations, confirming that the GFP-positive structures were the sites of active RNA synthesis, as they co-localized with EU that was incorporated into newly synthesized RNA, even when cellular transcription was blocked by actinomycin D, confirming that the GFP-positive structures were indeed VFs.

Next, we sought to understand more about the nature of IBDV VFs. We demonstrated that the VFs were not bound by a membrane, which is consistent with previous literature demonstrating that structures containing VP3 were visible following cellular permeabilization with digitonin (15). It has previously been documented that upon entering the cytoplasm, VP3 within the vRNP complex binds to phosphatidylinositol-3-phosphate on the cytoplasmic leaflet of endosomal membranes to seed the formation of the VF (16). Interestingly, the *Birnaviridae* are thought to form a phylogenetic link with single stranded positive (ss+) sense RNA viruses (1, 23), and the finding that the virus manipulates cellular membranes, presumably to provide a microenvironment more favorable for replication, is more akin to strategies used by ss+ RNA viruses in replication. Our finding that the VF is not bound by a membrane does not preclude the fact that there may be membrane structures within the VF, and Delgui *et al* have demonstrated endosomal membrane markers co-localize with VP3 –positive structures in QM7 cells infected with IBDV, even at 36 hpi (24). However, we did not observe any membrane structures within the VFs by TEM, raising the possibility that they are transiently present within a VF.

Given that the VFs were not bound by a membrane, we hypothesized that they could be formed by liquid-liquid phase separation (LLPS). This phenomenon is potentially a means of cytoplasmic compartmentalization without the need for membrane-bound organelles (25), and is known to be involved in the replication of other viruses, for example vesicular stomatitis virus, rotavirus, rabies virus, influenza A viruses, and measles virus (22, 26–29). Structures that form through LLPS typically show fusion and fission, rapid recovery of fluorescence after photobleaching, and are dissolved by 1,6-hexanediol (30, 31). We have previously demonstrated that IBDV VFs move rapidly in the cytoplasm, and coalesce together (17). Here, we extend these observations by demonstrating that VF FRAP was rapid, and the VFs were dissolved in the presence of 4% 1,6-hexanediol. Taken together, these data are consistent with the *Birnaviridae* VFs being formed through LLPS.

Next, we transfected cells with individual IBDV proteins tagged to GFP and observed that VP3-GFP formed VFLs, but that other IBDV proteins did not. Moreover, as VP1-GFP was ubiquitously present throughout the cytoplasm when expressed alone, we hypothesize that VP3 recruits VP1 to contribute to the formation of the VFs. This is consistent with other studies that have expressed VP3 alone in cells and found it forms distinct cytoplasmic puncta (16). Here we extend these observations by demonstrating that VP3 when expressed alone shows some FRAP, recovering to 46% of the original intensity over the course of the experiment, compared to VP4 which formed aggregates that showed no FRAP. This indicates that VP3 spontaneously forms cytoplasmic VFLs with some liquid-like properties, however, optimal LLPS may require the full complement of proteins and RNA to be in the VF.

We demonstrated that the VFs show a high level of co-localization with VP3 (Manders’ coefficient 0.9), but a lower co-localization with the VP2 capsid protein (MC 0.6). This observation is consistent with Dalton & Rodriguez (2014), who observed a lower colocalization of VP2 with VP1, VP3 or dsRNA at early time points post-infection (13). In order for this to be the case, the site of virus replication (the VFs) must be distinct from the PAs of virions that we and others have observed by TEM (32). In order to reconcile this, we conducted CLEM, and demonstrated that the VFs were discrete electron dense regions of the cytoplasm, separate from the PAs. However, key questions remain, including whether any cellular proteins or mRNAs are recruited to the VF, and what processes occur there. For example, it remains unknown whether translation occurs in the VF, or whether components of the innate immune pathway are sequestered in the VF, as they are in the inclusion bodies of respiratory syncytial virus (33). Moreover, whether there is an ultrastructure or compartmentalization within the VF remains unknown. While the TEM images show a homogeneous electron dense region, it is possible that proteins are not evenly distributed within the VF. Imaging using super-resolution microscopy would help to shed light on this. Finally, the dynamics of virion assembly and PA formation remains poorly understood. We did not observe VFs that contained visible virus particles by TEM, unlike avian reovirus. One hypothesis for this is that individual virus particles formed in the VF may be rapidly repelled into the cytoplasm due to a charge incompatibility. When multiple virions are expelled from a VF, they may accumulate together and form a lattice due to electrostatic interactions between the capsids. Alternatively, VP2 may accumulate in a VF until it reaches a critical concentration and then spontaneously forms many capsids, forming a PA. In this way, a PA may be a remnant of a VF. *Birnaviridae* are thought to undergo random packaging rather than selective packaging (34), and while both these models would be consistent with random packaging, much work is left to do in order to determine the process of virion assembly.

Our study is not without limitations. First, the GFP signal was only visible from 8 hpi using our equipment. Second, it has been shown that rotavirus VFs become less liquid over time (22). We only conducted FRAP and 1,6-hexanediol treatment at 16 hpi, and we did not compare different time-points. However, that FRAP was rapid and VFs dissolved by hexanediol at this time point suggests that the VFs do not lose fluidity over time in the same way as rotavirus VFs. Third, we conducted our studies within DF-cells which are immortalized fibroblast cells, whereas the tropism of IBDV *in vivo* is in B cells. It is challenging to image B cells, but owing to their more spherical nature, large nucleus, and their non-adherence it is possible that the IBDV VFs within the cytoplasm have different dynamics.

In summary, *Birnaviridae* VFs are sites of *de novo* RNA synthesis, are not bound by a membrane, show properties consistent with liquid-liquid phase separation, and are distinct from the PAs observed by TEM. This new information enhances our fundamental knowledge of the replication of a family of viruses responsible for disease of economic importance to poultry and aquaculture, that could be used to develop strategies to better control disease, or optimize their therapeutic application in the future.

## Acknowledgements

We are grateful for financial support from the BBSRC (grants BBS/E/I/00001845, BBS/E/I00007034, and BBS/E/I/00007039). The funders had no role in study design, data collection and interpretation, or the decision to submit the work for publication. In addition, we are grateful to Jόse Castόn of Centro Nacional de Biotecnologia (CNB-CSIC) for the anti-VP4 antibody.

## Figure Legends

**Movie S1. IBDV virus factories were present in infected, mock-treated cells imaged live.** DF-1 cells were transfected with GFP1-10 and infected with the PBG98-VP1-GFP11 virus. Infected cells were mock-treated with media 16hpi and imaged live over a time course of 17 minutes. Fifty z-stacks were imaged in total, with each stack being acquired at an interval of 20 seconds.

**Movie S2. IBDV virus factories dissolved in infected cells treated with 4% 1,6-hexanediol and imaged live.** DF-1 cells were transfected with GFP1-10 and infected with the PBG98-VP1-GFP11 virus. Infected cells were treated with a solution of 4% 1,6-hexanediol 16hpi and imaged live over a time course of 17 minutes. Fifty z-stacks were imaged in total, with each stack being acquired at an interval of 20 seconds.

**Movie S3. Paracrystalline arrays of virions detected by electron tomography.** DF-1 cells were infected with IBDV strain PBG98 and prepared for TEM 18hpi. Serial electron microscopy datasets were collected every 1^⍛^ over a 120^⍛^ tilt series and reconstructed. 10nm gold antibodies acted as fiducial markers.

## Notes

### Competing Interest Statement

The authors have declared no competing interest.

